# Substance P receptor blockade is associated with increased survival of patients with *C. difficile* infection

**DOI:** 10.64898/2026.09.06.749186

**Authors:** Leo Zhao, Min Liu, Siyu Wang, Melissa A. Musser, Amy Shepherd, Jie Zhang, Michael Whalen, David Binion, Meenakshi Rao, Min Dong, John Manion

## Abstract

*Clostridioides difficile* is a major cause of healthcare associated infections. *C. difficile* infection (CDI) is a toxin-mediated disease driven by the secretion of two large protein toxins, Toxin A (TcdA) and Toxin B (TcdB). TcdA and TcdB cause neurogenic inflammation that can contribute to intestinal inflammation in rat and mouse models. Substance P (SP), a potent vasoactive neuropeptide, is a major contributor to C. difficile toxin induced inflammation in animal models. SP acts on multiple receptors including the high affinity receptor NK1-R, and lower affinity receptors (*Mrgprb2* and *NK2-R)*. Through *in vivo* mouse infection models, we confirmed the effects of SP in hypervirulent *C. difficile* are largely mediated by the neurokinin-1 receptor (NK1-R), the high affinity receptor for SP. Through genetic mice that lack mast cells (cKit^W-sh^) we determined that mast cells which respond to SP in certain kinds of neurogenic inflammation appear dispensable for inflammation caused by *C. difficile* infection. We explored the role of NK1-R antagonists in a retrospective study of a large clinical cohort. We determined that, after propensity matching, NK1-R antagonists were associated with a significant survival benefit and that patients treated with NK1-R antagonists showed evidence of reduced vascular inflammation (increased serum albumin). Together these data suggest blocking SP activity through antagonism of NK1-R with small molecule inhibitors may be a useful strategy for adjunctive treatment during *C. difficile* infection.

## Introduction

*Clostridioides difficile* infections cause severe healthcare-associated infections that are associated with high mortality rates^1^. *C. difficile* occurs in the context of a disrupted colonic microbiota, through factors that include cancer treatment, inflammatory bowel disease and antibiotic use^2-5^. Recurrence and severe disease remain significant clinical challenges^6^. While antibiotic therapy with vancomycin or fidaxomicin can treat *C. difficile*, therapeutic options to manage severe fulminant infections remain limited^7^.

*C. difficile* is a toxin mediated disease and secretion of Toxins A and B (TcdA and TcdB) is required for pathology^8,9^. TcdA and TcdB promote inflammation when administered within the intestine, in skin tissues^10,11^ or when administered systemically^12^. Consistently, inflammation severity is associated with severe outcomes in *C. difficile*. Consequently, serum albumin, circulating cytokines and leukocyte counts are biomarkers for *C. difficile* severity^13^.

Immune responses are essential for bacterial clearance, however recent studies suggest targeting inflammation through modulation of immune responses may be an effective strategy for treating *C. difficile* infections. Several studies have also connected toxin mediated inflammation to colonization and disease^14,15^. Recent work also reinforces that vascular inflammation is an essential component of *C. difficile* infection^16^. Toxin A has long been associated with neurogenic vascular inflammation^17,18^ while recent studies have determined this is also the case for Toxin B^10^. *C. difficile* toxins intoxicate cells through receptor mediated endocytosis, and delivery of a toxic enzymatic glucosyltransferase cargo to the causing inactivation of Rho-GTPases^19^. Recent studies using single cell, genetic mice and fluorescent *in situ* hybridization indicate the *C. difficile* toxin receptors are enriched on specific cell-types within the intestine rather than being broadly distributed^10,20^. Toxin receptors for all major forms of TcdB are highly expressed on pericytes – specialized cells that wrap endothelial cells (CSPG4), while toxin variants also target intestinal stem cells in colonic crypts^21^ (Frizzled) and intestinal neurons (Frizzled and Tissue Factor Pathway Inhibitor-TFPI1^22^). Pericytes and neurons are consequently highly sensitive to the actions of TcdB^10,23,24^. At low doses of toxins, likely to occur early during infections, the action of the toxins on the neurons innervating the GI tract and local vasculature appears to be enhancement of neuronal signaling including neuropeptide secretion^10,23^, and cytokine secretion^10,16^. Of these neuropeptides, substance P (SP, encoded by *Tac1*), a vasoactive neuropeptide, is a major contributor through actions on the neurokinin-1 receptor (NK1-R) that contributes to edema and inflammation.

In mouse models of both infection and intestinal intoxication, NK1-R appears to be a major contributor to pathology through enhancement of inflammation^10,18^. These effects are conserved between both TcdA and TcdB. However, the role of NK1-R in human CDI remains unknown. A single prior study observed that infected human colons showed evidence of enhancement of NK1-R signaling during *C. difficile* colitis^17^. Other prior clinical studies of NK1-R antagonists suggested these drugs might lead to neutropenia in pediatric bone cancer patients^25^, which could possibly worsen *C. difficile* infection^26^. Consequently, while NK1-R antagonism appears beneficial in murine models^10^, the impact of NK1-R antagonists in human CDI remains undefined.

NK1-R antagonists are widely used for the prevention and treatment of nausea and emesis in patients undergoing chemotherapy with highly emetogenic chemotherapy agents. Cancer chemotherapy patients are at high risk of developing *C. difficile* through immunosuppression, microbiota disruption and impacts of chemotherapy on the colonic epithelium. Thus, we reasoned analysis of this population may reveal impacts of NK1-R antagonists on *C. difficile* outcomes.

## Results

### NK1-R knockout increases survival and significantly decreases pathology and colonization during infection with epidemic strains of *C. difficile*

Epidemic strains of *C. difficile* in the RT027 lineage frequently encode a variant of TcdB (referred to as TcdB2) which interacts with CSPG4^12,27^ but not Frizzled1, 2, and 7^27^; variants can also interact with alternate cellular receptors that include TFPI-1^22^ and Low-density lipoprotein receptor-related protein 1 (LRP1)^28^. TcdB2-producing strains also shows variation in processing domains^29^ and may differ with respect to inflammatory pathways and cellular targets^30^. We previously determined that treatment with NK1-R receptor antagonists is effective in both infections with TcdB1 and TcdB2, and that *Tac1*^*-/-*^ mice show increased protection from infections with strains that produce either toxin^10^. To extend these findings and determine whether *NK1-r* knockout is sufficient to protect from infection with epidemic strains, we infected mice with M7404^TcdA-TcdB+^, which expresses TcdB2 at high levels. Mice infected with M7404^TcdA-TcdB+^ produce high amounts of toxin^31^, show high mortality rates, and severe histopathological changes including edema (vascular inflammation), neutrophilic infiltrates and epithelial pathology^32^. Like *Tac1* deficient mice, and antagonist treated mice^10^, *NK1r*^*-/-*^ mice showed reduced colonization burdens (**Fig 1a**), a trend toward increased survival (**Fig 1b**), and significantly reduced histopathological damage to the intestine (**Fig 1c-g**). These data collectively further confirm that the actions of SP release following infection with hypervirulent strains including on pathology are largely mediated through NK1-R and that knockout of NK1-R reduces the colonization burden *of C. difficile* during infection.

**Figure 1:**
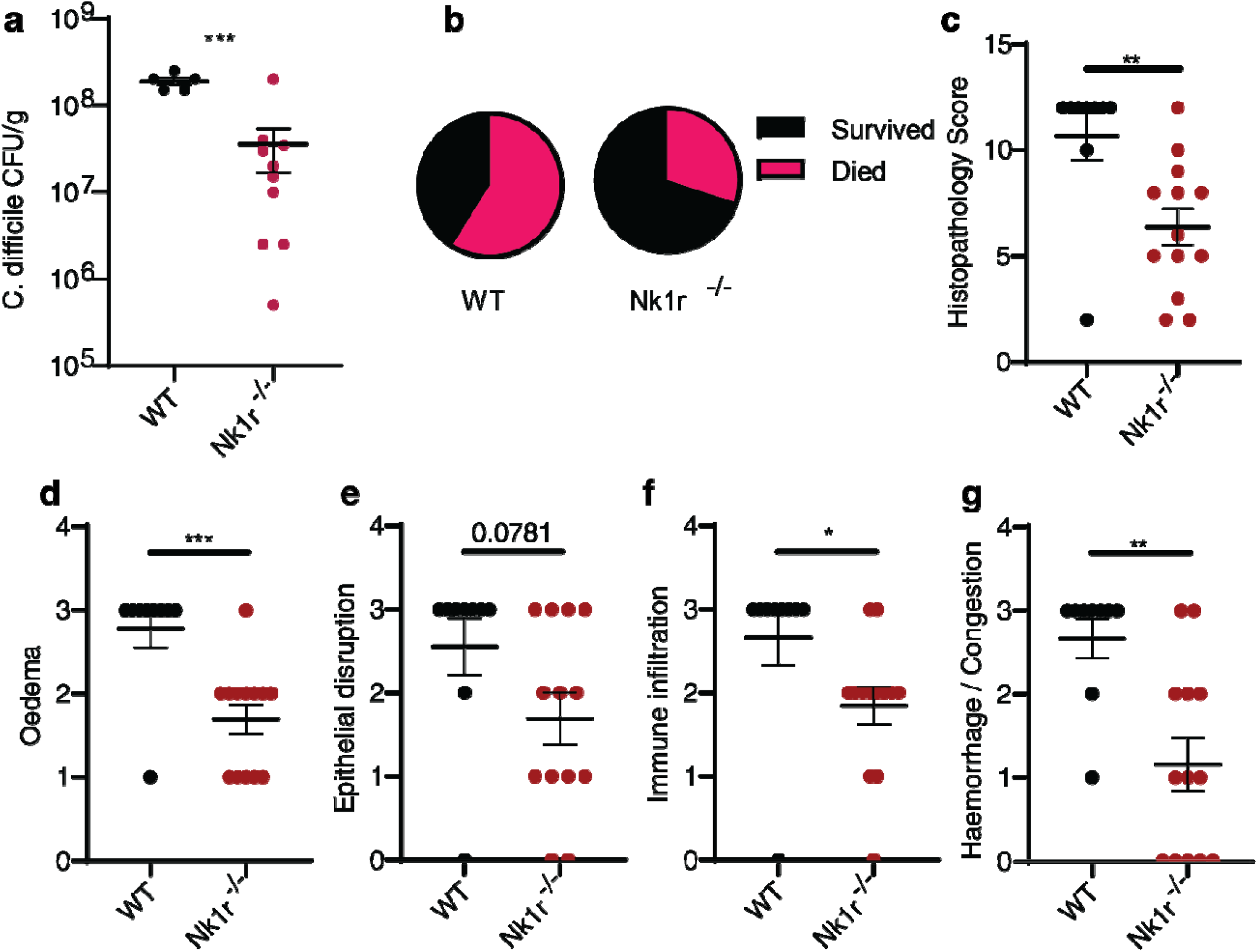
NK1R knockout protects against infection with M7404^TcdA-TcdB+^. (**a**) Mice were infected and colonization was quantified from cecal contents by colony forming units after 2 days. (**b**) Survival was assessed after two days of infection. N=17, WT and N=18 NK1r^-/-^ mice (**c**) Histopathology total scores were assessed for cecal pathology, sub scores are for (**d**) Oedema, (**e**) Epithelial disruption, **(f)** immune infiltration and **(g)** haemorrhage or congestion. P values are results from students’ two-tailed t-test. *<0.05, **<0.01, *** <0.001, means are presented and error bars reflect standard error of the mean.

### Mast cells are not required for CDI mediated pathology during infection in mice

Prior animal studies on *C. difficile* conducted using small intestine intoxication models with Toxin A suggested that mast cells may contribute to *C. difficile* toxin associated pathology^17^. Substance P is reported to function on two classes of receptor, Mrgprb2 and NK1-R. Prior studies have determined that, in certain cases, NK1-R antagonists can act on Mrgprb2 in mice but not humans (MrgprX2 in humans)^33^ which is suspected to be a challenge in clinical translation of NK1-R antagonists to humans^34^. Mrgprb2 is mainly localized to mast cells, and there are conflicting data about the expression of NK1-R in mast cells^35^. In skin, neurogenic inflammation can involve the integration of neuronal and mast cell responses, and these are reported to rely on *Mrgprb2*^*35*^. The localization of mast cells within the intestine and their impact on diseases varies^36^, so it is unknown whether a neuron-mast cell circuit contributes during infection in anatomically relevant large intestine tissues. Our previous results suggest skin inflammation elicited by TcdB is independent of mast cells^10^, but the potential role of mast cells in toxigenic infection in large intestine remains unknown. We infected mice that lack mast cells with *C. difficile* 630^Δerm^, a laboratory adapted strain of *C. difficile* that produces both TcdB1 and TcdA. We found that *C-kit*^*swsh*^ mice had similar cecal histopathology to wildtype animals (**Fig 2b**), suggesting that the impacts of these toxins are independent of mast cells in the cecum. Notably there were no changes in epithelial disruption, haemorrhage or congestion, inflammatory infiltrates or oedematous inflammation indicating these processes are independent of mast cells in this model (**Fig 2c-2f**). Under these conditions, we were unable to detect a substantial impact of mast cells on CDI suggesting that mast cells do not play a substantial role in protection from inflammation offered by *Tac1* knockout, *Nk1r* knockout or NK1-R antagonists. These data provide a further rationale for clinical application of NK1-R antagonists, since these drugs only antagonize NK1-R in humans^34^.

**Figure 2:**
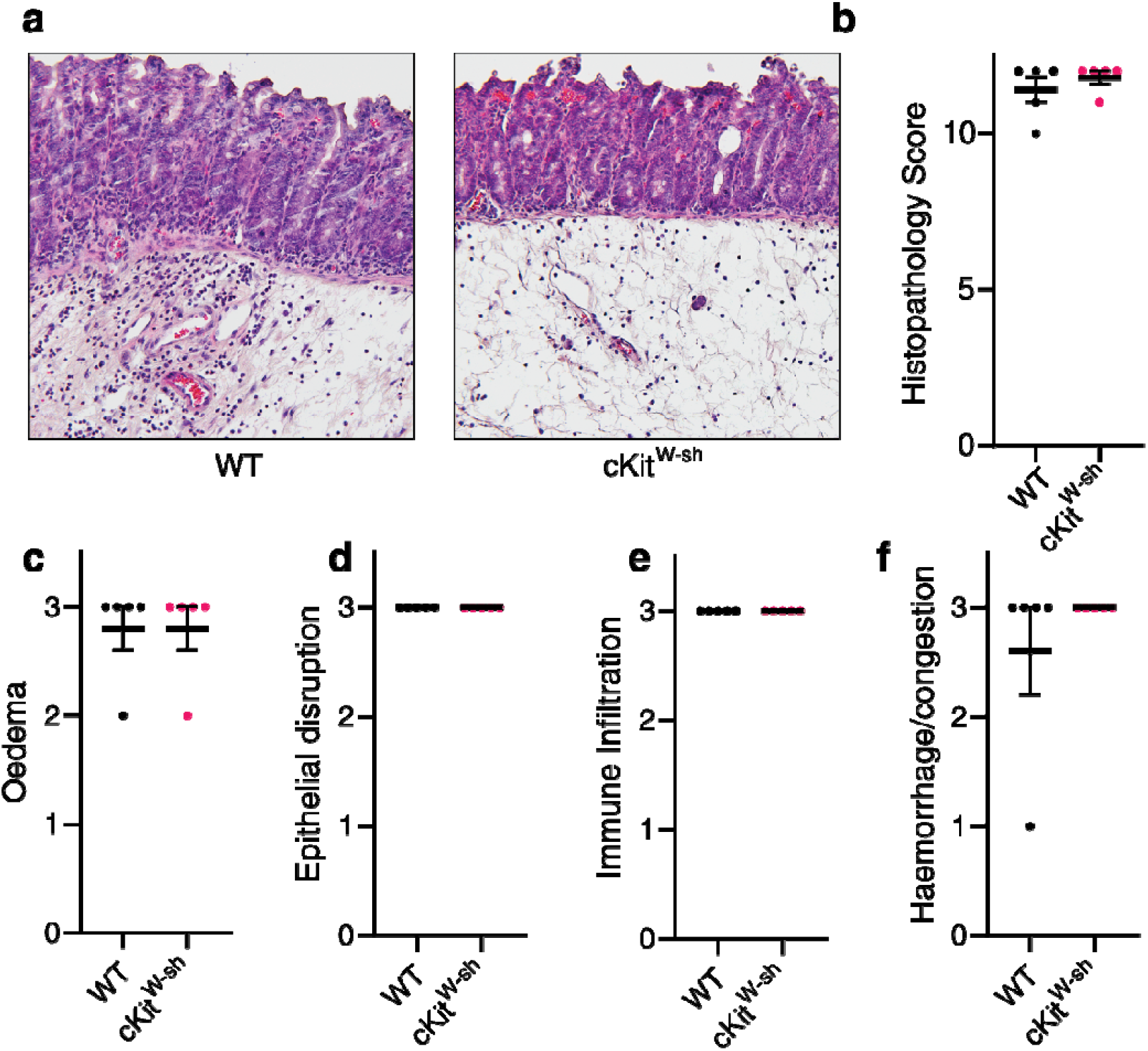
Mast cells are dispensable for *C. difficile* induced inflammation. Mast cell deficient cKit^W-sh^ and wild-type mice were infected with C. difficile 630, after two days histopathology was assessed to assess impacts on infection inflammation. (**a**) Representative micrographs showing similar inflammation quantified in **(b)** histopathology scores and sub scores are for (**d**) Oedema, (**e**) Epithelial disruption, **(f)** immune infiltration and **(g)** haemorrhage or congestion. Means are presented and error bars reflect standard error of the mean. No statistical significance was detected using a student’s two tailed t-test n=5 mice per group.

### NK1-R antagonists are associated with increased survival during CDI in humans

To expand our prior studies on the impacts of NK1-R blockade into human infection. We conducted a retrospective study to assess impacts of NK1-R antagonists (aprepitant, fosaprepitant and netupitant) on CDI outcomes using the TriNetX database^37^. We extracted *C. difficile* cases with NK1-R antagonist prescription within a 30-day period preceding and including the index date. Propensity matching was used to generate a control cohort with similar demographic factors, comorbidities, and chemotherapeutics; all matched characteristics have SMD<0.1 (**Table 1)**. Following propensity matching, we assessed 30-day mortality in 5,539 matched patients. NK1-R antagonists were associated with a statistically significant reduction in *C. difficile*-associated 30-day mortality (p=2.2 x 10^-6^, **Fig 3a**), with no overlap in 95% CI from days 16-30 post infection.

**Table 1:** Baseline and post propensity score matched cohort characteristics. SD Standard deviation, PSM, Propensity score matching. Matching was assessed by standardized mean difference.

| Variable |  | Aprepitant | Controls | Aprepitant (post-PSM) | Controls (post-PSM) | SMD post |
| --- | --- | --- | --- | --- | --- | --- |
| **Cohort size** |  | 5,539 | 429,846 | 5,536 | 5,536 |  |
| **Age at Index (years)** | Mean (SD) | 54.98 (19.82) | 60.09 (21.78) | 55.00 (19.80) | 55.97 (22.94) | 0.045 |
|  | Min - Max | 0 - 90 | 0 - 90 | 0 - 90 | 0 - 90 |  |
| **Sex** | Female | 3,161 (57.1) | 244,834 (57.0) | 3,160 (57.1) | 3,137 (56.7) | 0.008 |
|  | Male | 2,375 (42.9) | 184,784 (43.0) | 2,375 (42.9) | 2,397 (43.3) | 0.008 |
| **Race** | White | 4,050 (73.1) | 305,895 (71.2) | 4,048 (73.1) | 4,029 (72.8) | 0.008 |
|  | Black or African American | 585 (10.6) | 49,544 (11.5) | 584 (10.6) | 558 (10.1) | 0.015 |
|  | Asian | 271 (4.9) | 11,237 (2.6) | 271 (4.9) | 255 (4.6) | 0.014 |
|  | Unknown | 367 (6.6) | 45,380 (10.6) | 367 (6.6) | 428 (7.7) | 0.043 |
| **Ethnicity** | Hispanic or Latino | 414 (7.5) | 26,874 (6.2) | 414 (7.5) | 427 (7.7) | 0.009 |
|  | Not Hispanic or Latino | 4,523 (81.7) | 298,852 (69.5) | 4,521 (81.7) | 4,410 (79.7) | 0.051 |
| **Comorbidities n (%)** | Neoplasms | 5,353 (96.6) | 167,060 (38.9) | 5,350 (96.6) | 5,360 (96.8) | 0.01 |
|  | Endocrine | 5,176 (93.5) | 349,932 (81.4) | 5,173 (93.4) | 5,191 (93.8) | 0.013 |
|  | Digestive system | 5,112 (92.3) | 343,884 (80.0) | 5,110 (92.3) | 5,136 (92.8) | 0.018 |
|  | Blood/immune | 5,021 (90.7) | 260,748 (60.7) | 5,018 (90.6) | 5,001 (90.3) | 0.01 |
|  | Circulatory system | 4,629 (83.6) | 330,296 (76.8) | 4,629 (83.6) | 4,741 (85.6) | 0.056 |
|  | Injury, poisoning | 4,216 (76.1) | 250,102 (58.2) | 4,215 (76.1) | 4,298 (77.6) | 0.036 |
|  | Nervous system | 4,183 (75.5) | 261,775 (60.9) | 4,182 (75.5) | 4,274 (77.2) | 0.039 |
|  | Musculoskeletal | 4,119 (74.4) | 286,176 (66.6) | 4,119 (74.4) | 4,236 (76.5) | 0.049 |
|  | Genitourinary system | 4,083 (73.7) | 302,511 (70.4) | 4,083 (73.8) | 4,174 (75.4) | 0.038 |
|  | Respiratory system | 3,971 (71.7) | 285,159 (66.3) | 3,970 (71.7) | 4,129 (74.6) | 0.065 |
|  | Mental, behavioral | 3,308 (59.7) | 244,031 (56.8) | 3,307 (59.7) | 3,442 (62.2) | 0.05 |
|  | Skin/subcutaneous | 2,808 (50.7) | 198,858 (46.3) | 2,808 (50.7) | 3,041 (54.9) | 0.084 |
|  | External causes | 2,005 (36.2) | 148,046 (34.4) | 2,005 (36.2) | 2,117 (38.2) | 0.042 |
|  | Symptoms/signs | 5,476 (98.9) | 392,077 (91.2) | 5,473 (98.9) | 5,466 (98.7) | 0.012 |
|  | Factors influencing health | 4,755 (85.8) | 339,532 (79.0) | 4,752 (85.8) | 4,808 (86.8) | 0.029 |
| **Antineoplastic n (%)** | Antineoplastic, other | 4,318 (78.0) | 33,916 (7.9) | 4,315 (77.9) | 4,593 (83.0) | 0.127 |
|  | Alkylating agents | 2,097 (37.9) | 9,575 (2.2) | 2,095 (37.8) | 1,849 (33.4) | 0.093 |
|  | Antimetabolites | 1,936 (35.0) | 23,672 (5.5) | 1,933 (34.9) | 1,989 (35.9) | 0.021 |
|  | Antineoplastic antibiotics | 1,520 (27.4) | 6,290 (1.5) | 1,518 (27.4) | 1,300 (23.5) | 0.09 |

**Figure 3:**
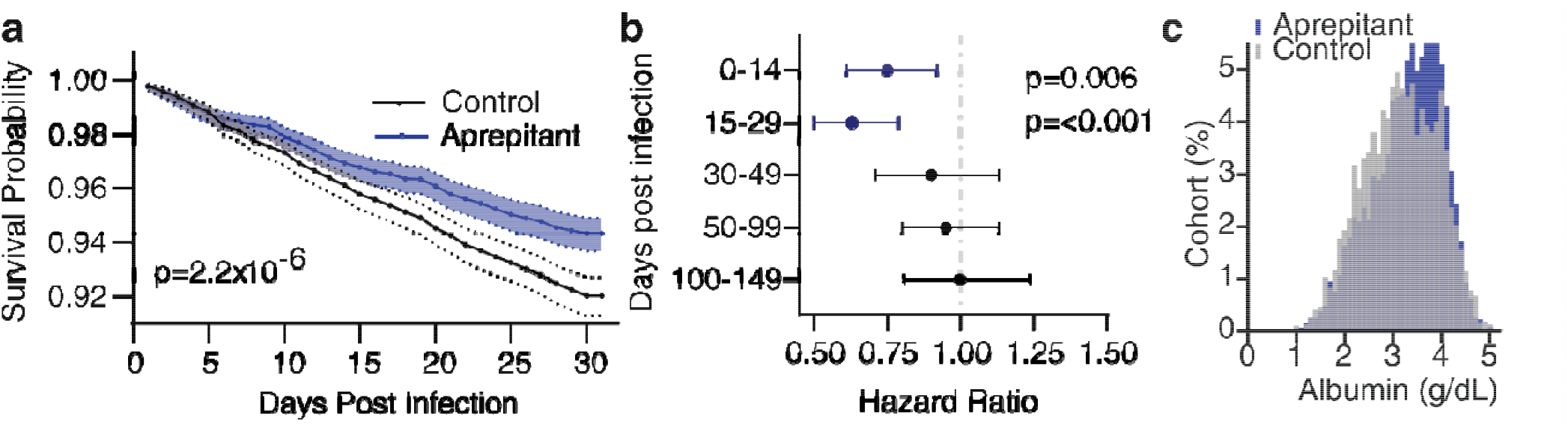
SP Antagonists (Aprepitant) use is associated with reduced risk of *C. difficile* associated mortality in a large multinational retrospective study. (**a**) Kaplan-Meier plot of survival in propensity matched cohorts (**table 1**) in the 30 days following a *C. difficile* infection. (**b**) Hazard ratios plotted overtime after index event (*C. difficile* infection). A lower hazard ratio indicates higher survival in aprepitant treated group. (**c**) Frequency distribution of albumin concentrations in the blood of *C. difficile* infected patients within 30 days of infection illustrating that albumin concentrations are higher in SP antagonist treated patients (table 2). Aprepitant group also includes other less frequently used NK1-R antagonists (see methods). Log rank test was used in panel a. See full statistical methods for panel **b**. Error bars reflect 95% confidence intervals.

Alternative hypotheses include that NK1-R antagonists improve all cause survival independently of CDI or the matched control cohort had a greater propensity for allcause mortality. If NK1-R antagonists improve all cause survival or the matched cohort were sicker, we would expect a sustained survival benefit in the NK1-R antagonist cohort for a prolonged period. To account for these possibilities, we assessed the hazard ratio over time stratified intervals up to 150 days post CDI. We observed a significant reduction in mortality post-CDI infection from 0-14 days (HR 0.75, 95% CI 0.61-0.92, p=0.006) and 15-29 days (HR 0.63, 95% CI 0.50-0.79, p<0.001). Beyond the 30-day interval, mortality benefits in the NK1-R antagonist cohort were no longer sustained (30-49, 50-99, and 100-150 days; all p>0.35). Together these data suggest reduced mortality in the NK1-R antagonist cohort following CDI were confined to the 30 days following infection (**Fig. 3b**).

**Table 2:** Laboratory findings in *C. difficile* cases post propensity matching, treated with either aprepitant or controls.

| Lab / Category | Aprepitant | Control | p-value |
| --- | --- | --- | --- |
| **Leukocytes (WBC, 10<sup>3</sup>/μL)** |  |  |  |
| 0–3 | 954 (20.7%) | 959 (24.3%) | <0.001 |
| 4–14 | 3,197 (69.5%) | 2,636 (66.9%) | 0.012 |
| 15+ | 451 (9.8%) | 344 (8.7%) | 0.091 |
| n | 4,602 | 3,939 |  |
| **ESR (mm/h)** |  |  |  |
| < 30 | 89 (36.5%) | 105 (37.8%) | 0.760 |
| 30 – <100 | 137 (56.1%) | 146 (52.5%) | 0.406 |
| ≥ 100 | 18 (7.4%) | 27 (9.7%) | 0.343 |
| n | 244 | 278 |  |
| **CRP (mg/L)** |  |  |  |
| < 10 (normal) | 273 (50.4%) | 225 (44.2%) | 0.045 |
| ≥ 10 (elevated) | 269 (49.6%) | 284 (55.8%) | 0.045 |
| n | 542 | 509 |  |
| **Albumin (g/dL)** |  |  |  |
| n | 4,732 | 3,840 |  |
| Mean | 3.29 | 3.16 | <0.001 |
| SD | 0.72 | 0.77 |  |

Symptomatic *C. difficile* causes a protein-losing enteropathy that is characterized by a loss of serum albumin and accumulation of blood proteins in feces^38^. Previously it was established that NK1-R antagonists reduce the intensity of inflammation caused by *C. difficile* toxins during infection^10,17,18^, especially edematous inflammation that reflects plasma protein extravasation caused by vascular inflammation. To assess indirect impacts on vascular inflammation, we examined blood test results within 30 days of a *C. difficile* infection to examine potential effects of NK1-R antagonists on serum-protein losing enteropathy. NK1-R antagonists were associated with increased serum albumin (**Fig 3c, Table 2**), suggesting reduced frequency of hypoalbuminemia. These data suggest that suppressing NK1-R activity may reduce loss of serum albumin from plasma protein extravasation during human CDI.

*C. difficile* infections are associated with increased inflammation, including elevated white blood cell counts and increased in inflammatory markers. We examined blood markers for inflammation to assess the impact of NK1-R antagonists. NK1-R were associated with lower proportion of patients with elevated C-reactive protein (49.6% vs 55.8%, p=0.045). ESR did not differ between cohorts (**Table 2**). Interestingly, given prior reports of neutropenia in patients receiving these drugs^25^, we found no association between NK1-R antagonists and increased risk of leukopenia; treated patients were less frequently leukopenic in this cohort (20.7% vs 24.3%, p<0.001) (**Table 2**). Collectively these data imply NK1-R antagonism may be beneficial during CDI by reducing mortality and decreasing the extent of serum albumin loss during infection without compromising immune cell numbers.

## Discussion

In a large clinical dataset, our results suggest that NK1R antagonism is associated with reduced mortality during CDI. Moreover, those data suggest NK1-R antagonism is associated with reduced loss of serum albumin during *C. difficile* infection. This data supports the hypothesis that NK1-R antagonism may improve survival in part through reduction of vascular inflammation which is reported to be an essential component of *C. difficile*-induced inflammation.

Prior studies identified potential roles of mast cell mediated effects of *C. difficile*^*25*^. Substance P is known to exert effects on mast cells, though recent studies suggest that many of these impacts are through expression of *Mrgprb2*^*33-35*^. We were unable to identify any major impact of mast cells on inflammation during CDI in mast cell deficient mice, suggesting that in this model of severe infection, mast cells are likely to only have a limited role. This is consistent with our genetic findings and prior pharmacological studies that most of the effects are through NK1-R^39,40^, which may not be expressed in mast cells. The role of mast cells varies by location through the intestinal tract, expression levels of NK1-R in cecal mast cells or the model of disease as these factors might impact the observed role of mast cells^36^. NK1-R knockout mice show strong but incomplete protection from infection, emphasizing the complex nature of *C. difficile* induced inflammation which includes (at least) epithelial^21^, immune^13^, neural^10,41^ and vascular components^16^.

Our retrospective studies provide support for the hypothesis that NK1-R antagonism is protective in humans. There are many limitations to retrospective designs, including selection bias, limited data availability and uncontrolled confounders. These studies largely include patients receiving NK1-R antagonists during chemotherapy. It remains unclear if these results will be the same in patients acquiring *C. difficile* through other causes such as inflammatory bowel disease, or immunosuppression. Moreover, many of these patients would also be receiving other antiemetics such as dexamethasone (corticosteroids) and ondansetron (5-HT3 antagonists) as part of three drug combinations used for chemotherapy induced nausea^42^. We also cannot rule out the possible impacts of patient selection such as physicians withholding unnecessary medications during *C. difficile* infections in severely unwell patients which could alter these results. Direct prospective randomized placebo-controlled trials of the role of NK1-R antagonists as adjuncts to antibiotic therapy would be necessary to rule out these potential effects.

Classically, approaches to bacterial infection and other host-pathogen interactions have focused on direct targeting of pathogens. However, these approaches have limitations that include antigenic variation, escape from antibodies, and evolution or adaptation of bacterial resistance. Increasingly, new approaches have suggested targeting the host to treat infections. This might be a useful approach to antibiotic resistance. While the host receptors targeted by *C. difficile* toxins are different, their final pathological effects, and cellular mechanisms appear largely conserved. Overall, antagonism of SP signaling through NK1-R offers a therapeutic opportunity for treating *C. difficile* infection.

## Author contributions

**Conception:** JM, LZ, DB, MR, MD **Study design:** LZ, MM, AS, MR, MD **TriNetX analysis**, LZ, MW, DB, JM **Clinical Advice:** LZ, MM, MR, MW, DB **Animal Studies** ML, LZ, MM, AS, SW, JZ, JM, MR and MD. **Supervision**, MW, MR, MD, JM. **Funding acquisition:** MR, MD, JM. **Manuscript original draft**, LZ, JM. **Editing:** All authors.

**Acknowledgements**

We thank all members of the Manion, Rao and Dong laboratory for technical assistance and comments. We thank Seth Rakoff-Nahoum for use of his anerobic chamber.

## Funding

This research was funded in part by the University of Pittsburgh School of Medicine. Work on *C. difficile* has been supported in part by The Assistant Secretary of War for Health Affairs endorsed by the Department of War through the Peer Reviewed Medical Research Program under Award Number HT9425-24-1-0047 to J.M. Opinions, interpretations, conclusions, and recommendations are those of the author(s) and are not necessarily endorsed by the Department of War. This research was supported in part by the National Institutes of Health, R01DK135707 to M.R., R01AI139087 to M.D. as well as core facilities provided by the Harvard Digestive Diseases Center (P30DK034854 from NIH)

## Data availability

TriNetX is a federated data source, data are readily available to collaborating institutions but cannot be publicly shared. Details of the data extraction are provided in the methods.

## Conflicts of Interest

The authors declare this work was conducted in the absence of financial conflicts of interest.

## Figures

## Methods Animal Studies

*Nk1r* ^*-/-*^ (Tacr1-CreERT2 mice)^43^, and congenic cKit^w-sh^ were obtained from Jackson Laboratory. *NK1r* ^*-/-*^ mice were compared to C57BL6/N mice, and fully congenic cKit^w-sh^ were compared to C57BL6/J mice as controls. 6–14-week-old male and female mice were used for these experiments, and sexually dimorphic impacts were not observed. Animal experiments were conducted at Boston Children’s Hospital under a protocol (1465) approved by the institutional animal care and use committee. Experiments took place in a vivarium which was temperature and humidity controlled, animals were fed a diet of normal chow and water ad libitum. Animals were housed in individually ventilated cages and bred under specific pathogen free conditions. Infection experiments took place in an BSL-2 environment which contains mouse specific pathogens. Euthanasia was by CO2 inhalation followed by cervical dislocation. Sample sizes were determined based on author experience with these models rather than formal power calculation.

### Spore preparation

Spores of 630Δerm, M7404^TcdA-TcdB+^ (Bacteria were gifts of Dena Lyras)^8,32^ were prepared as previously described. Cultures were grown in a sporulation broth for 5 days; cells were collected by centrifugation and washed twice in sterile PBS before being shocked with 50% v/v ethanol at room temperature for 1 hour. The suspension was washed five times in PBS before collecting spores. Spores were stored at -80C and quantified on *C. difficile* ChromID agar plates (Biomerieux) after 24 hours in an anaerobic chamber.

### CFU enumeration

*C. difficile* were isolated from cecal content, weighed and dissolved in pre-reduced phosphate buffered saline (PBS). Cecal contents were diluted and enumerated on ChromID *C. difficile* agar plates as previously described^10^. Live bacteria were counted by performing multiple dilutions and assessed after 24 hours of incubation in an anerobic chamber (Coy Labs).

### Histopathology

Following euthanasia by CO2 inhalation, cecal tissues were collected, cecal content was removed for CFU analysis, then ceca were flushed with PBS before being transferred to 10% neutral buffered formalin for fixation. Tissues were processed and embedded into paraffin by the Beth Israel Deaconess Medical Center (BIDMC) core facility. Blocks were subsequently sectioned at 6µM before being stained by hematoxylin and eosin. Slides were baked, deparaffinized then rehydrated through a series of ethanol washes. Slides were immersed in Gill’s hematoxylin (No3, Thermo Fisher, Shandon) prior to washing with Scott’s tap water (0.1% sodium bicarbonate) to blue, prior to immersion in acidic eosin. Slides were dehydrated in serial changes of ethanol and xylene before being mounted in DPX. Sections were imaged and scored blindly according to our published criteria briefly; these are composed of 0-3 scores (0- none, 1- mild, 2-moderate, 3-severe), for oedema, epithelial disruption, immune infiltration and vascular haemorrhage or congestion^10^. Only animals that survived the full observation period were included in histopathology scoring.

### Infection models

Mice were infected with *C. difficile* through a previously established protocol.^16^ All infections were performed in an A-BSL2 facility with autoclaved cages, chow and water. Mice were fed with an antibiotic cocktail containing vancomycin (0.4mg/mL), colistin (850 U/mL), metronidazole (0.215mg/mL), gentamicin (0.035 mg/mL) and kanamycin (0.045mg/mL) diluted in drinking water for 3 days. This was switched to regular autoclaved water for 2 more days then mice were injected with clindamycin (10mg/kg) by intraperitoneal injection. 24 hours later mice were gavaged with spores 10^4^ of 630^Δerm^, M7404^TcdA-TcdB+^ as indicated. Mice were monitored twice a day and euthanized if they became moribund (trouble moving or breathing, or >15% body weight loss).

### Clinical data source

This retrospective cohort study utilized the TriNetX platform, a federated global database of 172 healthcare organizations (HCOs) which provide de-identified electronic medical records of more than 205 million patients. Non-identifiable information is exempt and not categorized as human subject’s research by the University of Pittsburgh Institutional Review Board (determination under STUDY2026-9-27966). The HCOs are located across North and South America, Europe, the Middle East, Africa, and Asia-Pacific. The platform contains comprehensive patient-level records which include but not limited to demographics, diagnoses, procedures, medications, lab values, vital signs, and genomic information. All data usage is compliant with the Health Insurance Portability and Accountability Act (HIPAA, § 164.514 (b) (1)), EU General Data Protection Regulation (GDPR, Regulation 2016/679), and ISO 27001:2022 certification. Analytics in the TriNetX platform are conducted at the HCO level - only aggregated de-identified results are returned to the database. Individual patient records do not leave contributing HCOs and HCO identities are not disclosed to prevent re-identification.

### Cohort selection

*C. difficile*-associated enterocolitis patients were identified using International Classification of Diseases, 10^th^ Revision, Clinical Modification (ICD-10-CM) code A04.7. Using PubChem Compound Identifier, patients receiving aprepitant, netupitant and fosaprepitant (CID: 358255, 1552337, 1731071) were classified as the NK1RA cohort, while those who never received these medications were classified as NK1RA non-receivers. A total of 435,385 patients with C. difficile-associated enterocolitis were identified, comprising 5,539 NK1RA-treated patients and 429,846 NK1RA-untreated patients.

### Assessment of covariates

Propensity score matching (PSM) was performed on 4 demographic factors, 15 comorbidities, and 4 classes of chemotherapeutics to balance the treated and untreated cohorts and minimize confounding. For demographic factors, age at index date, sex, race, and ethnicity were matched. Comorbidities were identified using ICD-10 diagnostic code ranges and selected based on both clinical significance as potential confounders and statistically significant differences between the two groups in pre-matching analysis. Comorbidities included diseases of the digestive, circulatory, respiratory, genitourinary, musculoskeletal, and nervous systems, as well as neoplasms, endocrine and metabolic disorders, diseases of the skin and subcutaneous tissue, diseases of blood and blood-forming organs, injury and poisoning, external causes of morbidity, mental and behavioral disorders, and factors influencing health status. NK1RA are commonly given with chemotherapy and different types of chemotherapeutic regimen may independently affect survival outcomes. Therefore, chemotherapeutics were identified using PubChem Compound Identifiers and selected as covariates to minimize confounding. These include antineoplastic alkylating agents, antineoplastic antimetabolites, antineoplastic antibiotics, and other antineoplastic agents.

### Assessment of outcomes

The index event was defined as the date of the first recorded diagnosis of *C. difficile* enterocolitis. NK1RA use was required to be within a 30-day period preceding and including the index date. After PSM, clinical outcomes in the 30 days following the index event were assessed and compared between NK1RA-treated and untreated cohorts. Outcomes assessed included survival and lab markers of systemic inflammation. Laboratory outcomes included serum albumin (g/dL), leukocyte count (10^3^/uL), C-reactive protein (mg/dL), and erythrocyte sedimentation rate (mm/h).

### Statistical analysis for clinical data

Baseline characteristics of treated and untreated cohorts were compared before and after PSM. For continuous variables (age), values are presented as mean ± standard deviation and compared using a two-tailed t-test. For categorical variables (sex, race, ethnicity, comorbidity, and treatments), values are presented as frequencies and percentages and compared using chi-squared test.

Cohort balance was evaluated using the standardized mean difference (SMD), with SMD < 0.1 considered indicative of adequate balance. Propensity score matching was performed in a 1:1 ratio, matching on demographic factors, comorbidities, and chemotherapeutics. Propensity score matching and assessment of cohort balance were performed using TriNetX platform’s analysis features.

Survival analysis of the groups over 30 days following *C. difficile* enterocolitis was performed using the Kaplan-Meier method with the log-rank test (**Fig 3a**). Cox proportional hazards regression analysis was performed to compare treatment effects across 150-day follow-up window using patient data reconstructed from TriNetX reported Kaplan-Meier survival probabilities, baseline cohort size, and total event counts^44,45^. The hazard ratio of the reconstructed cohort matched the TriNetX-reported overall hazard ratio to within 2%. To test if the hazard ratio is constant over time, we performed Schoenfeld test. Schoenfeld test was significant (p=0.006), indicating hazard ratio varied over follow-up period. Therefore, we defined time intervals within the followup period (0-14, 15-29, 30-49, 50-99, and 100-150 days) and calculated separate hazard ratios for each time interval. Interval specific hazard ratios with 95% confidence intervals are displayed as a forest plot (**Fig 3b**). These analyses were performed in R version 4.5.1.

Distribution of albumin are visualized as relative frequency histograms for each cohort, with values grouped into clinically relevant intervals and y axis representing percentage of patients within each interval. Inflammatory markers C-reactive protein (CRP), erythrocyte sedimentation rate (ESR), and leukocyte count are compared between treated and untreated cohort. All three markers are presented as mean ± standard deviation and between-group differences were assessed using two-tailed t test.

### Statistical analysis for animal studies

Data were collated in Microsoft Excel; statistical analyses were performed in GraphPad Prism. Student’s Two-Tailed T test was used to compare between groups. Data are presented +/- Standard error of the mean

## Notes

### Competing Interest Statement

The authors have declared no competing interest.

